# Combinatorial mosaics dissect the residue level basis of ACE2 binding divergence between SARS-CoV-2 and RaTG13

**DOI:** 10.64898/2026.10.13.758899

**Authors:** Simone M. Giovanetti, Meru J. Sadhu

## Abstract

The spike proteins of SARS-CoV-2 and the closely related bat coronavirus RaTG13 differ in their binding affinity of human ACE2. This is due to differences in the receptor binding domain (RBD), the region primarily responsible for ACE2 binding, which are only 89% identical on the amino acid level, compared to 96% for the remainder of their genomes. Here, we constructed a combinatorial library of RBD mosaics containing combinations of 18 amino acid differences between SARS-CoV-2 and RaTG13 and measured human ACE2 binding using yeast surface display and fluorescence-activated cell sorting. The mosaics revealed a broad range of binding phenotypes. The effects on ACE2 binding were concentrated at eight positions, including all six variable positions that directly contact ACE2. Although the SARS-CoV-2 spike binds human ACE2 stronger than RaTG13 spike, the SARS-CoV-2 residues Q493 and Q498 each reduced ACE2 binding relative to RaTG13’s tyrosines. By measuring the effects of these residues across many combinations of other sequence differences, we identified sequence context dependence for residues 493 and 498, finding that the deleterious effects of Q493 and Q498 were attenuated by the SARS-CoV-2 residues at positions 483, 484, and 494. Together, these results provide a residue-level account of the sequence differences underlying the difference in ACE2 binding by SARS-CoV-2 and RaTG13 RBDs, including which differences have the largest effect and which depend on sequence context. This work demonstrates that mosaic libraries provide a robust approach for resolving the contributions of sequence differences across diverse backgrounds.

**Significance Statement:** Closely related proteins can differ greatly in function. Identifying which sequence differences underlie the difference in phenotype can be difficult. We built a large library of receptor binding domains that independently combined 18 differences between SARS-CoV-2 and the related bat coronavirus RaTG13 and measured their relative binding to human ACE2. Most of the difference in binding was concentrated to eight positions, and the effects of two of these positions, 493 and 498, changed depending on nearby amino acids. These results show that combinatorial mosaic libraries can resolve functional differences between homologous proteins at the residue level while also revealing context dependent effects.

## Introduction

SARS-CoV-2, the coronavirus responsible for the COVID-19 pandemic, relies on the spike glycoprotein to enter host cells. The receptor-binding domain (RBD) of spike initiates this process by binding ACE2 on the host cell surface (1). During the COVID-19 pandemic, mutations in spike repeatedly altered viral infectivity, immune evasion, and protein stability (2–4). The ability of spike to bind ACE2 is an important determinant of host range and its emergence as a human pathogen (5,6).

RaTG13, a bat coronavirus sampled from *Rhinolophus affinis*, is a close relative of SARS-CoV-2 (5,7). Across the whole genome, they share 96% amino acid identity (5) but only 89% amino-acid similarity in the RBD (8). Most of the sequence differences distinguishing the two spikes cluster in the receptor-binding motif (RBM), a loop within the RBD that mediates receptor binding (Fig. 1A, 1B) (9). The SARS-CoV-2 RBM has 17 residues that directly contact ACE2, six of which are diverged between SARS-CoV-2 and RaTG13 (10). The RBMs also differ at 11 sites that do not directly contact ACE2. The two RBDs differ in their affinity for human ACE2, with RaTG13 spike binding human ACE2 approximately 70-fold less efficiently than SARS-CoV-2 (reported Kd 21.1 nM for SARS-CoV-2 vs 3.86 μM for RaTG13, by surface plasmon resonance) (10). This functional divergence raises the question: which of the naturally occurring sequence differences between SARS-CoV-2 and RaTG13 RBDs result in the different affinities for human ACE2?

**Figure 1.**
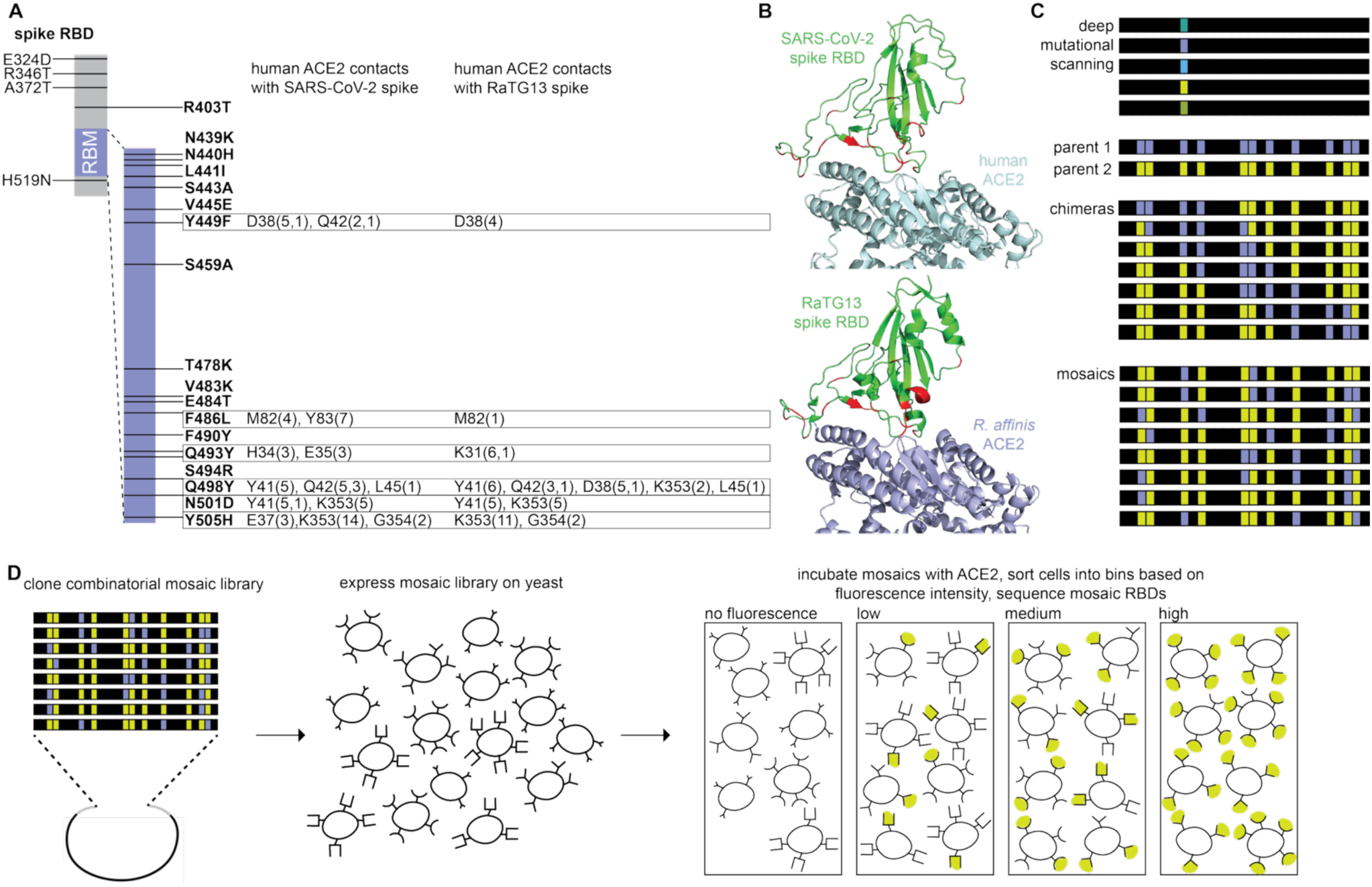
Design and functional characterization of a combinatorial SARS-CoV-2/RaTG13 RBD mosaic library. Sequence and structural comparisons of the SARS-CoV-2 and RaTG13 differences incorporated in the combinatorial mosaic library and assayed by yeast surface display and FACS. **A.** Amino acid differences between the SARS-CoV-2 and RaTG13 spike receptor binding domains (RBDs). Seventeen differences are within the receptor binding motif (RBM). Contacts made by these variable residues in either RBD to human ACE2, as determined from structural analyses (PDB 6LZG and 7DRV), are indicated (10). Numbers in parentheses indicate first the number of van der Waals contacts and second potential hydrogen bonds with the listed ACE2 residue (10). **B.** Crystal structure of SARS-CoV-2 spike RBD bound to human ACE2 (top; PDB 6M17) and RaTG13 RBD bound to *R. affinis* ACE2 (bottom; PDB 8Z9I). Identical sites between SARS-CoV-2 and RaTG13 are shown in green, while divergent sites are colored in red. **C.** Schematic comparison of genetic space sampled by deep mutational scanning, traditional chimeras, and combinatorial mosaics. Mosaic libraries independently assort the naturally occurring amino acid differences between two parental genotypes, allowing each residue to be sampled across many combinations of the other sequence differences. **D.** Experimental workflow for measuring ACE2 binding across the mosaic library. Mosaic RBD sequences were cloned into a yeast-display vector and expressed on the yeast surface. Cells were incubated with human ACE2 and, following fluorescent labeling of bound ACE2, sorted by fluorescence activated cell sorting into four bins based on fluorescence intensity (proxy for ACE2 binding). The abundances of mosaic RBD sequences in each bin were then quantified by high throughput sequencing to estimate the relative ACE2 binding of each genotype.

Several studies have identified candidate residues that contribute to this difference. Deep mutational scanning (DMS) studies comprehensively measured the effects of individual amino acid substitutions on ACE2 binding in the SARS-CoV-2 RBD. Substitutions corresponding to several RaTG13 residues reduced ACE2 binding, including those at positions 403, 449, 486, 501, and 505, whereas substitutions at positions 439, 440, 483, 493, 494, and 498 increased binding (11). Targeted mutagenesis studies examined the differences between the SARS-CoV-2 and RaTG13 RBD directly. The SARS-CoV-2 arginine at residue 403, which lies outside the RBM, enhances ACE2 binding relative to the RaTG13 threonine through direct electrostatic interaction with ACE2 residue E37 and stabilization of the spike trimer (12). Other studies focused on the 17 residues that differ between the RBMs of RaTG13 and SARS-CoV-2. In one study, six RBM residues in the RaTG13 spike were individually replaced with the SARS-CoV-2 residue. Four of these substitutions (F449Y, L486F, D501N, H505Y) enhanced ACE2 binding, while Y493Q reduced and Y498Q abolished binding. Simultaneous introduction of all six variants increased ACE2 binding, but the resulting construct still bound less strongly than wild-type SARS-CoV-2 RBD (10). A similar study found that RaTG13 RBD proteins with the SARS-CoV-2 amino acid at positions 449, 486, 493, 501, and 505 increased ACE2 usage in a cell-cell fusion assay, whereas K439N and Y498Q impaired function and T484E was neutral (13). Overall, the two studies reached largely concordant findings, aside from residue 493.

These studies illustrate a broader challenge in identifying the sequence determinants of functional divergence between homologous proteins. Most targeted mutagenesis studies assess substitutions in one or a small number of sequence backgrounds. Yet the effect of a substitution can depend strongly on the surrounding sequence. For example, the Q498R mutation that occurred in the Omicron lineage is deleterious in the original SARS-CoV-2 background but becomes strongly affinity-enhancing in the presence of an additional Omicron mutation, N501Y (14,15). More broadly, experiments performed with different SARS-CoV-2 spike variants have shown that the sequence context can reshape the effects of subsequently introduced mutations (15, 16). This context dependence complicates efforts to determine which sequence differences are determinants of phenotypic divergence.

A common approach to studying functional divergence between two homologous proteins is the use of chimeras, in which contiguous regions are exchanged between them. By characterizing a series of chimeras, investigators can localize regions that contribute to the phenotypic differences between the parental proteins. Chimeras have been extensively used to map functional domains across diverse protein families for decades (17–19). However, because each chimera replaces an entire sequence segment, causal residues cannot be distinguished from other sequence differences within the exchanged region. Furthermore, the relatively small number of constructs typically studied provides limited sampling of the sequence backgrounds in which each sequence difference can occur. Thus, traditional chimeras can localize the sequence determinants of functional divergence to the responsible regions but cannot resolve the contributions of individual residues or determine how those contributions vary across sequence backgrounds.

High-throughput DNA synthesis and sequencing have enabled the construction and characterization of large libraries of combinatorial variants in which naturally occurring sequence differences vary independently (16, 20–22). By breaking linkage between neighboring residues, these libraries repeatedly sample each parental residue across many combinations of the other naturally occurring differences (Fig. 1C). We refer to these variants as mosaics to distinguish them from traditional chimeras, reflecting the independent assortment of sequence differences rather than the exchange of contiguous regions. Mosaics can therefore be used to estimate which residues have effects that persist across diverse genetic backgrounds while also revealing those whose effects depend on sequence context.

Here, we applied this high-throughput mosaic approach to understand which naturally occurring differences between the SARS-CoV-2 and RaTG13 RBDs account for their divergent binding to human ACE2. We focused on 18 amino acid differences, including the 17 differences within the RBM and one immediately outside it. We generated a library of thousands of mosaic RBDs, each carrying a distinct combination of naturally occurring residues from the two parental sequences. By characterizing their ability to bind human ACE2 in high-throughput, we identified the residues that distinguish ACE2 binding phenotypes of SARS-CoV-2 and RaTG13 while revealing cases in which their effects depend on the surrounding sequence.

## Results

### Combinatorial mosaic library samples SARS-CoV-2 and RaTG13 differences across diverse genetic backgrounds

The SARS-CoV-2 and RaTG13 spike protein RBDs differ at 22 amino acid positions (8). We focused on the RBM (residues 438-506), the segment of the spike RBD that mediates ACE2 binding, because it contains sequence differences most likely to underlie the divergence in receptor binding (23). We also included residue 403, which is outside the RBM, as it is nonetheless proximal to ACE2 and was previously implicated in ACE2 binding (12). In total, we examined 18 of the 22 differences between the two RBDs (Fig. 1A). Six of these occur at residues that were crystallographically determined to directly contact ACE2 (449, 486, 493, 498, 501, 505; 10). The remaining differences lie outside the direct ACE2 binding interface and may influence receptor binding indirectly (Fig. 1B).

To measure the effects of these naturally occurring differences across combinatorial backgrounds, we designed a library of combinatorial RBD constructs, which we call mosaics. At each of the 18 variable positions, a mosaic carries either the SARS-CoV-2 or RaTG13 residue, generating 2^18^ (262,144) possible combinations (Fig. 1C). We leveraged oligonucleotide synthesis technology in which variable nucleotides can be incorporated at user-defined positions. For twelve variant residues that differ by a single nucleotide, both the SARS-CoV-2 and RaTG13 variant can be encoded in single oligonucleotides using variable nucleotide encoding. Thus, a set of 32 oligonucleotides of length 230 nucleotides encoded all 2^17 combinations of the possible RBM mosaics. Variation at residue 403 was introduced by cloning the oligonucleotide pool into parallel plasmid backbones encoding either the SARS-CoV-2 arginine or RaTG13 threonine at this position. This design breaks the linkage among the naturally occurring sequence differences, allowing each residue to be functionally assayed across many combinations of the other differences separating the two RBDs (Fig. 1D).

We generated a library of mosaic RBDs for yeast surface display and characterized its representation via PacBio sequencing. Of the 262,144 possible genotypes, 87,306 were present after processing filters, providing broad coverage of the combinatorial sequence space. Both parental amino acids were well represented at each variable position, with the frequency of the SARS-CoV-2 residue ranging from 40% to 59% across the 18 sites (Fig. 2A). Pairwise genotype frequencies closely matched those expected from the observed single site frequencies, with only minor deviations from the expected distributions (Fig. 2B,C). These results indicate that the combinatorial library was largely free of systematic biases and that there was effectively no linkage between positions, as pairwise combinations occurred close to their expected frequencies.

**Figure 2.**
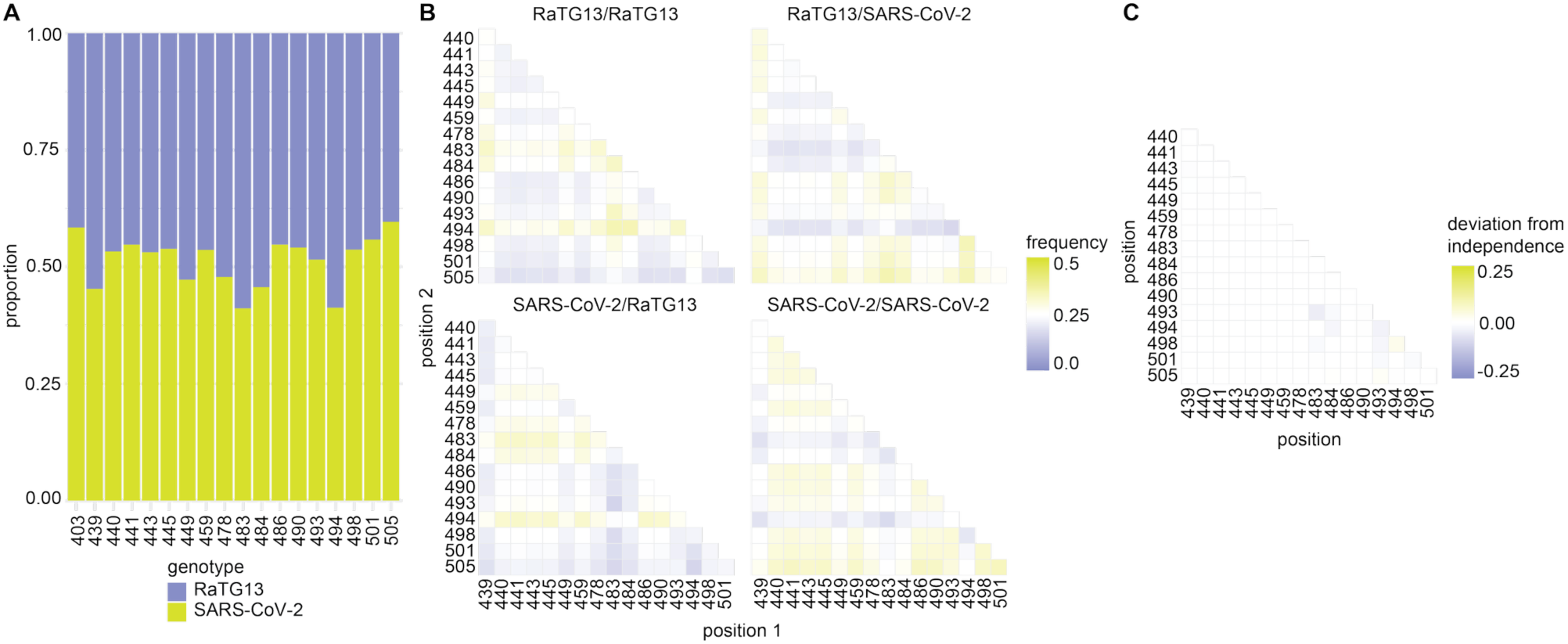
Representation and pairwise association of parental amino acids across the mosaic library. Parental amino acids were balanced across the mosaic library. **A.** Proportion of genotypes in Pacbio sequencing of the mosaic pool carrying the SARS-CoV-2 or RaTG13 amino acid at each of the 18 variable positions. **B.** Pairwise representation of parental amino acid combinations across the library. For each pair of variable positions, mosaics were classified into the four possible combinations (RaTG13/RaTG13, RaTG13/SARS-CoV-2, SARS-CoV-2/RaTG13, or SARS-CoV-2/SARS-CoV-2). Heatmaps show the observed frequency of each combination for every pair of positions. **C.** Deviation of observed pairwise representation from that expected under independent assortment of the two positions. Values near zero indicate that the pairwise combinations occurred close to the expected frequencies, whereas positive or negative values indicate over or underrepresentation of the mixed-parent combinations, respectively.

We measured human ACE2 binding across the pooled mosaic library using yeast surface display, fluorescence activated cell sorting (FACS), and sequencing (Fig. 1D). Yeast displaying the mosaic RBD library were incubated with 100 nM ACE2, a concentration at which binding by RaTG13 was weakly detectable. We then sorted the yeast into fluorescence bins spanning low to high ACE2 binding. We sequenced each bin to determine the distribution of individual mosaic genotypes across the fluorescence bins. To compare ACE2 binding between genotypes, we calculated a mean bin score from a genotype’s normalized abundance across the fluorescence bins, with higher scores indicating greater ACE2 binding (Fig. 1D).

### ACE2 binding effects are concentrated among eight RBD differences

Consistent with previous studies, the RaTG13 RBD exhibited substantially lower binding to human ACE2 than the SARS-CoV-2 spike (Fig. 3A). Recombining the sequence differences between the two RBDs generated a broad distribution of binding phenotypes that extended beyond both parental sequences. Among mosaic genotypes, 14% bound ACE2 more weakly than RaTG13, 59% had an intermediate phenotype between the two parental RBDs, and 26% exceeded the binding observed for SARS-CoV-2 (Fig. 3A). Thus, different combinations of the naturally occurring sequence differences separating RaTG13 and SARS-CoV-2 generate a wide range of ACE2 binding phenotypes.

**Figure 3.**
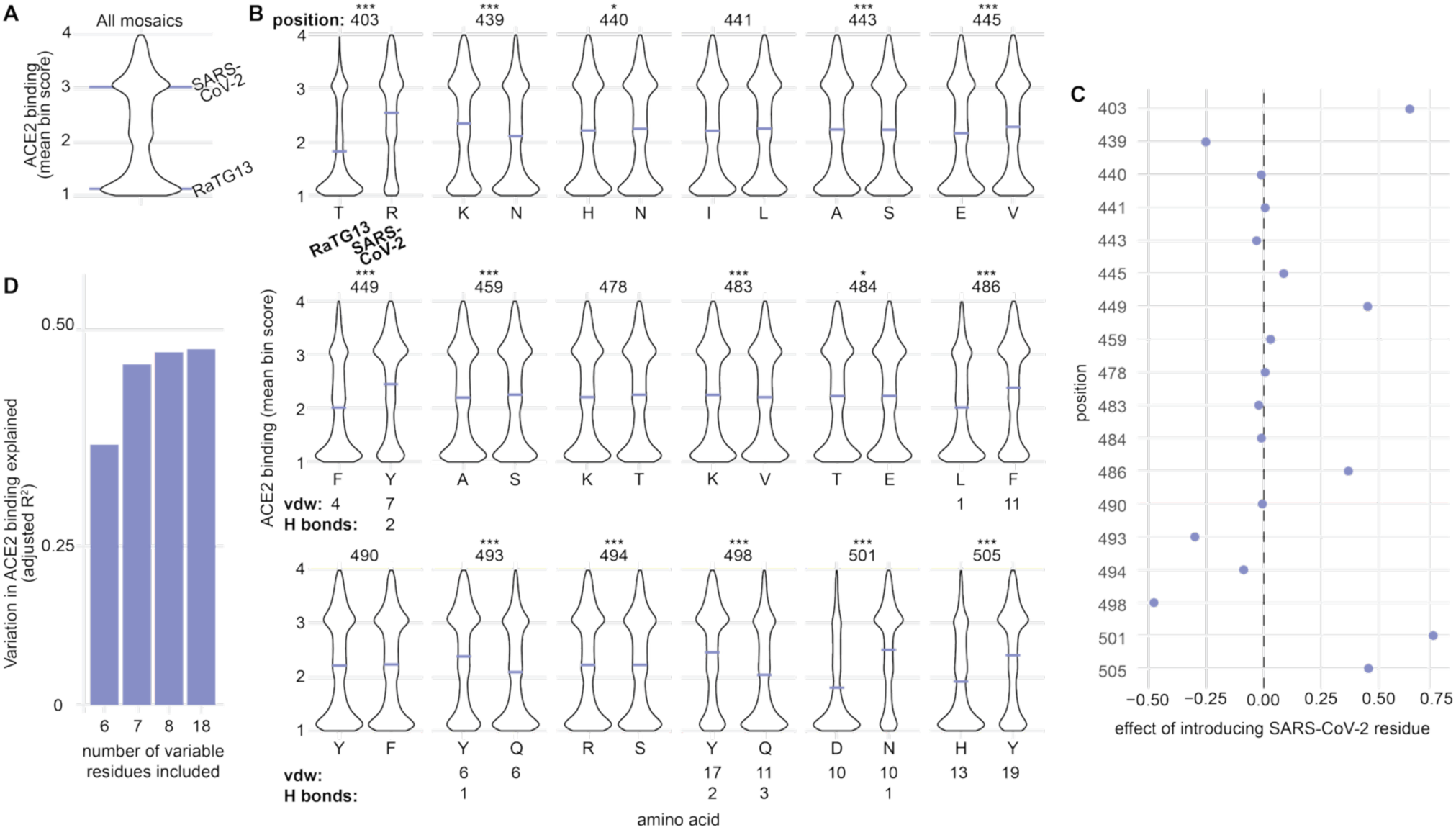
ACE2 binding effects are concentrated among a subset of variable residues. ACE2 binding varied broadly across the mosaic library with eight positions capturing nearly all of the variation explained by all 18 variable sites. **A.** Distribution of ACE2 binding phenotypes across the mosaic library. Relative binding was quantified as mean bin score, a weighted average, for each genotype. Horizontal lines indicate the mean bin scores for wild-type RaTG13 and SARS-CoV-2. **B.** ACE2 binding distributions according to amino acid identity at each of the 18 variable positions. For each position, mosaics were separated according to whether they carried the RaTG13 or SARS-CoV-2 residue, while averaging over variation at the remaining positions. Horizontal lines indicate the average mean bin score for each group. Statistical significance was determined from the coefficient for each position in an additive linear model containing all 18 positions. P values were adjusted for multiple comparisons using the Benjamini-Hochberg false discovery rate method (FDR < 0.05; adjusted p < 0.05 *, p < 0.01 **, p < 0.001 ***). Numbers under ACE2 contacting positions indicate the number of van der Waals contacts and potential hydrogen bonds formed with human ACE2 reported by Liu et al. (10). **C.** Average effect of introducing the SARS-CoV-2 residue at each position, calculated as the difference in mean bin scores between mosaics carrying the SARS-CoV-2 and RaTG13 amino acid at that position. Positive values indicate that the SARS-CoV-2 residue increased ACE2 binding on average. The dashed vertical line indicates no effect. **D.** Variation in ACE2 binding explained by additive linear models containing increasing numbers of variable positions, shown as adjusted R2. Models contain six variable ACE2 contact residues (449, 486, 493, 498, 501, 505), seven (+403), eight (+439), or all 18 variable positions.

We asked which of the 18 sequence differences contributed most strongly to variation in ACE2 binding. For each position, we partitioned the full panel of mosaics according to whether they had the SARS-CoV-2 or RaTG13 amino acid at that position and compared ACE2 binding between the two groups (Fig. 3B). Because each residue was sampled across many combinations of the other 17 variable positions, these comparisons estimate the average effect of each difference across many sequence backgrounds rather than in a single parental background.

The largest differences in ACE2 binding were associated with residues 403, 439, 449, 486, 493, 498, 501, and 505 (Fig. 3C). Mosaics carrying the SARS-CoV-2 residue at positions 403, 449, 486, 501, and 505 bound ACE2 more strongly on average than those carrying the corresponding RaTG13 residue. In contrast, the SARS-CoV-2 residues at positions 439, 493, and 498 were associated with lower ACE2 binding on average. The remaining 10 sequence differences had comparatively small or undetectable effects across the mosaic library (Fig. 3B, 3C). The direction of effect at each position was broadly consistent with previous studies, including measurements performed in the Wuhan-Hu-1 SARS-CoV-2 background. These results validate the mosaic framework and demonstrate that the assay recapitulates known determinants of spike-ACE2 interactions.

Six of the eight positions with the largest effect (449, 486, 493, 498, 501, 505) were previously defined to be on the ACE2 binding surface (10). In many cases, the favorable SARS-CoV-2 residues form stronger or additional contacts with ACE2 than their RaTG13 counterparts. For instance, SARS-CoV-2 residue Y449 makes more extensive van der Waals contacts and potential hydrogen bonds with ACE2 D38 and Q42, F486 forms additional van der Waals interactions with M82 and Y83, N501 forms a hydrogen bond with Y41 that is absent in RaTG13, and Y505 makes additional van der Waals interactions with K353 and a new interaction with E37 (10; Fig. 1A). Of the residues not defined as part of the binding surface, residue 403 lies outside of the RBM but has previously been implicated in receptor binding (12). In SARS-CoV-2, the N439K substitution, which introduces the RaTG13 residue, increases human ACE2 affinity by creating a salt bridge with E39 in ACE2 (24). Thus, all the positions with the largest average effects directly influence receptor binding.

We asked how much of the variation in ACE2 binding could be attributed to these positions using a variant component analysis. Main effects for the six previously characterized ACE2-contact residues in the RBM (449, 486, 493, 498, 501, 505) explained 37% of the variance in ACE2 binding across the combinatorial landscape, compared to 49% explained by all 18 positions. Adding residues 403 and 439 increased the variance explained to 49% (Fig. 3D). Thus, these eight positions captured nearly all of the variation explained by the 18 single position effects.

Although these eight positions had the largest average effect, their contributions varied across sequence backgrounds. We further probed the importance of sequence context, focusing on positions 493 and 498 because the SARS-CoV-2 residues at both sites were associated with reduced ACE2 binding despite occurring in the more strongly binding SARS-CoV-2 RBD. We therefore asked whether the effects of these variants differed across sequence backgrounds in the RBD.

### The effects of variants at residues 493 and 498 depend on sequence background

To determine whether the effects of variants at positions 493 and 498 depended on other sequence differences, we examined each position in combination with each of the other 17 variable positions. For each pair, we partitioned the mosaic library into the four possible combinations of the SARS-CoV-2 and RaTG13 residues at the two positions (Fig. 4A, C). This allowed us to compare the effect of changing residue 493 or 498 when the second position carried either parental amino acid, while averaging over the remaining sequence differences. Because mean bin scores are a monotonic but nonlinear proxy for ACE2 binding, we focused on interactions that changed the direction of a residue’s effect. Specifically, we identified cases in which the effect of Q493 or Q498 changed sign depending on the residue at a second position, or in which Q493 or Q498 changed the sign of a second position’s effect.

**Figure 4.**
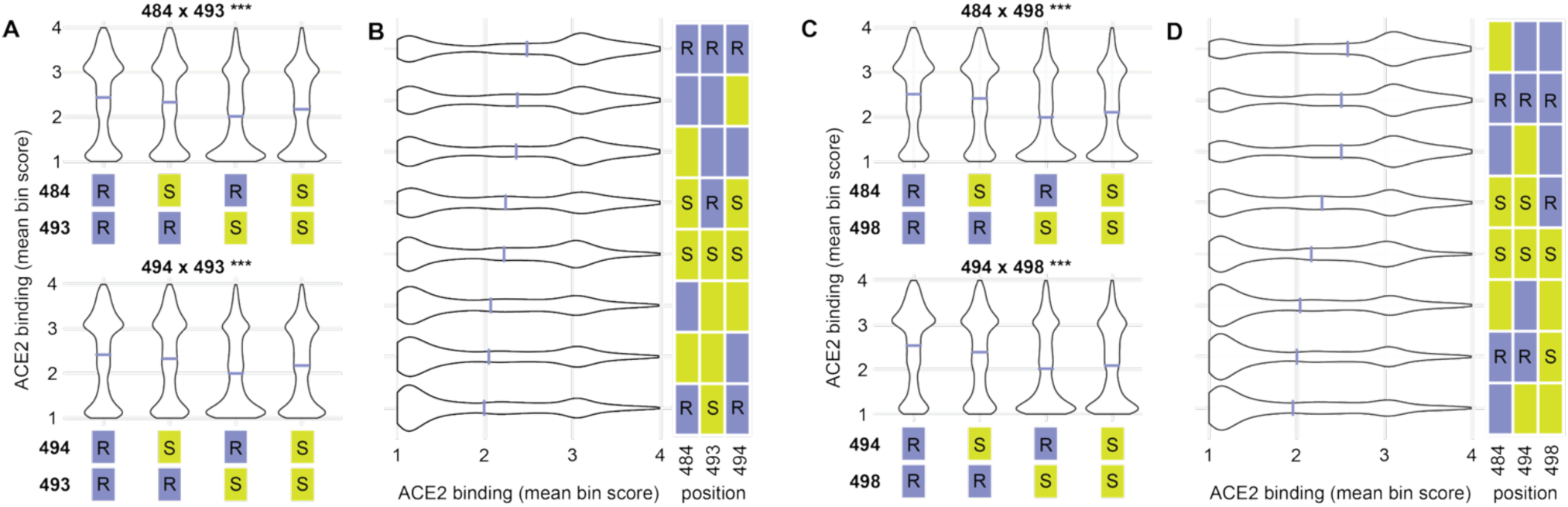
The effects of residues 493 and 498 depend on the sequence context. Pairwise and combined analyses identify sequence contexts that modify the effects of the SARS-CoV-2 residues Q493 and Q498. **A.** Pairwise analysis of residues 493 with positions 484 and 494. For each pair, mosaics were partitioned into four possible combinations of RaTG13 (R) and SARS-CoV-2 (S) amino acids at the two positions. Horizontal lines indicate the mean bin score for each group. **B.** Combined analysis of positions 484, 493, and 494. Mosaics were partitioned into all eight combinations of the parental amino acid states at these three positions. Violin plots show the distribution of ACE2 binding for each position and the adjacent matrix indicates the corresponding genotype (RaTG13 is purple, SARS-CoV-2 is yellow). Selected combinations are labeled to highlight the effect of changing 493 in otherwise all-RaTG13 or all-SARS-CoV-2 like contexts. **C.** Pairwise analysis of residue 498 with positions 484 and 494. D. Combined analysis of positions 484, 494, and 498. Mosaics were partitioned into all eight combinations, with genotypes indicated in the adjacent matrix. Selected combinations are labeled to highlight the effect of changing 498 in otherwise all-RaTG13 or all-SARS-CoV-2-like contexts. Statistical significance was determined from the interaction coefficient in a linear model containing the main effects at all 18 positions and all pairwise interactions among the 18 positions. For panels A and C, the indicated significance corresponds to the interaction term between the 493 or 498 and the indicated second position. P values for the interaction terms were adjusted for multiple comparisons using the Benjamini-Hochberg false discovery rate (FDR < 0.05; adjusted p < 0.001 ***)

We found sign-changing interactions between position 493 (Y in RaTG13 and Q in SARS-CoV-2) and positions 484 and 494. For instance, in the presence of Y493, mosaics carrying the RaTG13 residue T484 had stronger binding than mosaics with E484, while the opposite was true in the presence of Q493 (Fig. 4A). We observed similar effects at positions 494 (Fig. 4A).

We next examined the combined effects of 484 and 494 on 493 by partitioning mosaics according to all eight combinations of the parental amino acids at positions 484, 493, and 494 (Fig. 4B). In mosaics with RaTG13-like T484 and R494, we found that Q493 was substantially worse than Y493: the all-RaTG13 combination had the highest ACE2 binding, while replacing Y493 with Q493 in this context was the worst. In contrast, in the SARS-CoV-2-like E484 and S494 context, the effect of Q493 was mild. The combination containing Y493 ranked fourth, while the combination containing Q493, ranked fifth and was the highest-binding combination that included Q493.

Residue 498 (Y in RaTG13 and Q in SARS-CoV-2) showed a similar context dependence on positions 484 and 494 (Fig. 4C). In the presence of Q498, the SARS-CoV-2-like residues at 484 and 494 were superior, whereas in the presence of Y498, the RaTG13-like residues were superior (Fig. 4C). We therefore examined all eight combinations of the parental amino acids at positions 484, 494, and 498. In the RaTG13-like T484 and R494 context, the combination containing Y498 had the second-highest ACE2 binding, while Q498 reduced binding to the second-worst. In the SARS-CoV-2-like E484 and S494 context, the difference between Y498 and Q498 was smaller, with Y498 only slightly more favorable (Fig. 4D). The mosaic combination with all SARS-CoV-2 residues (E484, S494, Q498) was the highest-binding combination containing Q498.

These results show that examining variants across combinations of sequence backgrounds can reveal context-dependent effects. Although Q493 and Q498 reduced ACE2 binding on average across the mosaic library, the magnitude of these effects depended on the identities at other positions. Q493 and Q498 were better tolerated in the presence of E484 and S494. Thus, the functional consequence of a variant can depend on the sequence background in which it is measured.

### Combinatorial mosaic library identifies major effect positions with limited genotype sampling

The analysis here used more than 80,000 mosaics, but libraries of this size might not be practical in other applications. To explore the robustness of our results, we downsampled the number of analyzed mosaic genotypes. First, we asked how the number of sampled genotypes affected the estimated effect for each variant. We repeatedly downsampled the mosaic library and calculated the root mean square deviation of the resulting position effect estimates relative to those obtained from the complete library. We downsampled to subsets ranging from 100 to 50000 mosaics and repeated this process 1000 times for each subset size. The root mean square deviation from the full library decreased as the number of sampled genotypes increased (Fig. 5A). At 100 genotypes, the RMSD is 0.18, comparable to the effect we measured for site 439 (0.25), indicating the magnitudes of the effects would have substantial uncertainty. By 1000 genotypes, the RMSD dropped to 0.06, and estimates increasingly approached the full library values and became stable by approximately 10000 genotypes.

**Figure 5.**
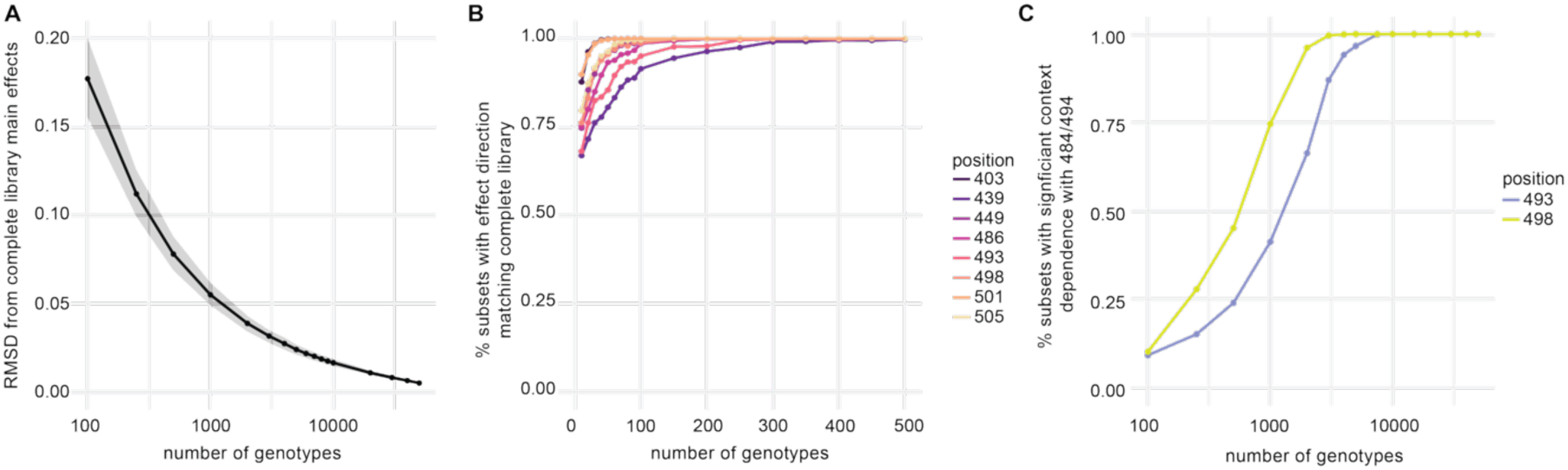
Major effects and sequence context dependence can be recovered without exhaustive sampling of the combinatorial mosaic library. Downsampling analyses assess the sampling required to recover average and context dependent effects. A. Effect of library size on the position effect estimates. The complete mosaic library was repeatedly downsampled across a range of genotype counts and, for each position, the difference in ACE2 binding phenotype between mosaics with the RaTG13 and SARS-CoV-2 residue was calculated for each subset. Points indicate the mean root mean square deviation (RMSD) between the 18 position effect estimates obtained from the downsampled library and the complete library. Gray halo indicates the interquartile range across 1000 replicate samples. **B.** Recovery of the direction of effect for the eight positions with the largest average effects. For each library size, the fraction of 1000 downsampled subsets for which the sign of the estimated effect matched that from the complete library is shown. **C.** Effect of library size on detection of context dependence at positions 493 and 498. Downsampled libraries were used to compare the effect of the RaTG13 and SARS-CoV-2 residue in two contrasting backgrounds (RaTG13- or SARS-CoV-2-like at 484 and 494). For each library size, the plot shows the fraction of 1000 downsampled subsets that, in a linear model, had a statistically significant difference for the effect of 493 and 498 in the contrasting backgrounds.

We separately asked how many genotypes were required to recover the direction of the effect for the eight positions with the largest average effects (403, 439, 449, 486, 493, 498, 501, 505). With 100 sampled genotypes, seven of the eight positions were assigned the correct direction in all 1000 replicate samples, while the remaining position (439) was correctly assigned 90% of the time. By 300 genotypes, the correct direction of effect was recovered for all residues 99% of the time (Fig. 5B). Thus, even a small panel of mosaics can reveal the variant sites with the strongest contributions.

Finally, we asked how library size affected our ability to detect the sequence context dependence observed at positions 493 and 498. For each downsampling, we looked at the effects of the residue of interest (493 or 498) in two contrasting backgrounds: the E484/S494 (SARS-CoV-2-like) or the T484/R494 (RaTG13-like) backgrounds. We fit a model testing whether the effect of 493 or 498 differed between the backgrounds and was statistically significant. Detection of context dependence increased with library size. At 1000 genotypes, we detected greater tolerance of Q493 in the SARS-CoV-2-like E484 and S494 background in approximately 40% of simulations; the corresponding context dependent effect of Q498 was detected in approximately 75% of simulations (Fig. 5C). By 10000 genotypes, these effects were detected 100% of the time. Detecting context dependent effects required broader genotype sampling than recovering the direction of the largest average effects, reflecting the greater sampling need to estimate the effect of a substitution within specific backgrounds.

By independently assorting differences from homologs across many sequence backgrounds, mosaic libraries therefore combine residue level information with an understanding of how consistent those effects are across a variety of sequence contexts. The amount of sampling required depends on the question being asked: major effects can be recovered from relatively smaller libraries, whereas more complex measures, like context dependence, require larger libraries.

## Discussion

### 493 and 498

We sought to identify which sequence differences between RaTG13 and SARS-CoV-2 RBDs are responsible for their divergent human ACE2 binding phenotypes. Our mosaic analysis showed that these effects were explained by eight positions and that the contributions of some positions depended on sequence context. This context dependence was particularly evident at positions 493 and 498. The effect of both residues 493 and 498 varied according to the identities of residues 484 and 494. The context dependence of residues Y493Q and Y498Q may reflect different configurations of the receptor-binding interface in RaTG13 and SARS-CoV-2.

At both positions, SARS-CoV-2 replaces a rigid aromatic tyrosine with a flexible polar glutamine. In RaTG13, Y493 hydrogen bonds with ACE2 K31, while Y498 has contacts with D38, Y41, Q42, and K353 (10). Introducing glutamine at either position into a RaTG13-like interface might remove favorable interactions made by the tyrosines. The positioning and orientation of those glutamines could, in turn, be affected by their environment. We found that the effects of Q493 and Q498 were ameliorated by E484 and S494. These two variants make similar changes to the RBM environment. At position 484, the glutamic acid in place of RaTG13’s threonine introduces a negatively charged side chain. The serine in place of RaTG13’s arginine at position 494 removes another positive charge. These residues therefore shift the RBM region toward a more negative electrostatic environment, which could alter the electrostatic and hydrogen bonding environment of the receptor binding ridge and influence the orientation or positioning of the glutamines, whose amide and carbonyl groups have opposite affinities for negatively charged surfaces, at the RBD-ACE2 interface. In addition, E484 lies near ACE2 K31, the same ACE2 residue contacted by Q493, providing another route by which changes at position 484 could influence the interactions made by Q493, and S494 can form an intramolecular hydrogen bond with Q493 (25), potentially helping stabilize a Q493 orientation compatible with interactions with ACE2 K31 and E35.

The effects of positions 493 and 498 suggest that the two RBDs may favor different local configurations of the receptor binding interface. The RaTG13 context better accommodates Y493 and Y498, while the SARS-CoV-2 context partially compensates for the less favorable Q493 and Q498 via changes in local electrostatics, hydrogen bonding, or geometry. Thus, the two RBDs may represent different but internally compatible configurations of the ACE2 binding interface, such that exchanging additional residues is required to compensate for the effect of the glutamines.

### Mosaic libraries complement single-site mutagenesis and chimera approaches

Our mosaic library approach complements both single-site mutagenesis approaches, like deep mutational scanning (DMS), and traditional chimera approaches. Single-site mutagenesis typically measures amino acid substitutions in a single sequence background, providing broad coverage of the amino acid effect but relatively limited sampling of the sequence context. In contrast, we restricted each variable position to the amino acids found in SARS-CoV-2 and RaTG13 but sampled those differences across many combinations of the other differences separating the two RBDs. For instance, Liu et al. (10) found that introducing all six SARS-CoV-2-like residues into the RaTG13 background did not fully restore SARS-CoV-2-like binding of ACE2, which could be due to missing the context of the additional variants that we explore here. Single-site mutagenesis asks how possible mutations alter a protein, mosaics ask how the differences between homologs contribute to functional divergence across combinations of those differences.

Mosaic libraries build on traditional chimeras. Exchanging contiguous regions between homologs can localize functional differences to particular regions but preserves linkage among the differences within the region. The resulting phenotype represents the combined effect of the variants in that window and cannot distinguish individual residue contributions. We observed this directly when we restricted our mosaic library to only consider genotypes containing a contiguous stretch of residues corresponding to SARS-CoV-2 within an otherwise RaTG13 backbone. In this reduced set of genotypes, the negative effects associated with the SARS-CoV-2 residues at positions 493 and 498 were not detected (SI Appendix, Fig. S1), because these residues remained linked to other variants with favorable effects on ACE2 binding. Thus, breaking linkage reveals individual effects that can be masked by other sequence differences contained within the same window. By independently assorting parental differences across many sequence backgrounds, mosaic libraries combine residue level understanding of their contribution to phenotype and the ability to determine how those effects are impacted by sequence context.

### Mosaic libraries do not require exhaustive sampling of the combinatorial space

A practical challenge of combinatorial approaches is that the number of possible genotypes increases exponentially with the number of variable positions. We therefore asked how much of the combinatorial space must be sampled to identify the residue differences that make the largest contributions to the difference in ACE2 binding by SARS-CoV-2 and RaTG13 RBDs. Our downsampling analysis suggests that exhaustive coverage is not required to recover the major determinants of functional divergence. Even with only 100 sampled genotypes, seven of the eight largest effect positions were assigned the correct direction of effect in every downsampled subset, and the eighth was correctly assigned in 90% of them. With 400 genotypes, the direction of all eight was recovered. These libraries represent a very small fraction of the 262,144 possible mosaics, indicating that sparse sampling can provide robust identification of the individual residues that most strongly dictate homolog specificity.

The amount of sampling required depended on the question being asked. While recovering the direction of the major effects required relatively few genotypes, estimating the magnitude required broader sampling, with estimates stabilizing for library sizes beyond 1000 genotypes. Smaller mosaic libraries can be used to identify candidate determinants of functional divergence, while more comprehensive sampling allows the estimation of the quantitative effect estimates or sequence context dependence.

Detecting sequence context dependence required additional sampling. We downsampled the library and asked how often we could recover the difference in the effects of residues 493 and 498 between the RaTG13-like T484/R494 context and the SARS-CoV-2-like E484/S494 context. Detection of these interactions increased with library size; by 3000 genotypes we could detect this interaction in more than 75% of downsamples for positions 493 and 498. This analysis demonstrates the difference in sampling depth required to detect the average effect of a residue versus whether that effect changes across sequence backgrounds. The latter requires sufficient representation of the residue of interest within the particular backgrounds and therefore benefits from greater sampling across the genotype space.

This scalability allows mosaic approaches to be extended to homologs separated by more differences than can be sampled exhaustively. Rather than constructing every possible combination, libraries can sample a subset of the genotype space in which each parental residue is represented across many backgrounds. This maintains the advantage of breaking linkage without requiring exhaustive sampling of the entire genotype space. The mosaic library size required will depend on the specific library assayed, its associated effect sizes, and the extent of context dependence, so the thresholds observed here are not universal. Nevertheless, our results show that exhaustive sampling is not required to resolve functional differences between homologs at the residue level. Even libraries representing a small fraction of the possible combinatorial space can robustly identify the major sequence determinants of functional divergence, while deeper sampling reveals dependence on sequence context.

### RaTG13

Among the sequence differences examined here, effects on human ACE2 binding were concentrated at eight positions, including six variable residues that directly contact ACE2. Most of the other differences between SARS-CoV-2 and RaTG13 RBDs had relatively little direct effect on receptor binding. This concentration of functional effects could have important consequences for RBD evolvability. If many positions can vary without substantially disrupting receptor binding, while changes at a small number of positions can strongly alter receptor binding, RBDs may be able to contain considerable sequence diversity while retaining the capacity to shift receptor affinity or specificity. Consistent with this, single amino acid substitutions in diverse sarbecovirus RBD backgrounds can alter the affinity or enable binding to novel ACE2 orthologs (15). A receptor binding surface that is determined by a small set of context-dependent residues could permit rapid shift in receptor specificity.

Conversely, the small effects of many RBD differences on human ACE2 binding suggest that these positions may vary in response to selection pressures other than receptor binding. Sequence variation outside the receptor binding determinants could affect antigenicity, folding, stability, spike conformation, or recognition of other ACE2 proteins. ACE2 itself is also variable. *Rhinolophus sinicus* populations contain multiple ACE2 alleles differing at residues within the RBD binding interface, and these alleles differ in their ability to bind different sarbecovirus spike proteins. Several of these interface residues show evidence of positive selection (16). More broadly, different sarbecovirus RBDs have different patterns of binding of *Rhinolophus* ACE2 proteins and some RBDs can use a broad range of ACE2 proteins (27). Thus, the functional consequence of RBD variation may depend on the ACE2 sequence.

An unresolved question is why the RaTG13 RBM is so divergent despite its close genomic relationship with SARS-CoV-2. Analyses suggest that much of this divergence occurred on the lineage leading to RaTG13. The highly divergent variable region containing the residues that contact ACE2 has been proposed to have been acquired by the RaTG13 lineage from an unsampled sarbecovirus through recombination (28, 29). In contrast, SARS-CoV-2 spike is highly similar to the spike protein from the BANAL coronaviruses isolated directly from bats (30), and thus the sequence differences examined here do not represent a series of substitutions through which SARS-CoV-2 progressively acquired human ACE2 binding.

What selective pressures produced the divergent RaTG13 RBM remain uncertain. One possibility is that the divergence occurred along the RaTG13 lineage to facilitate binding to a different host species’ ACE2. RaTG13 was isolated from an *R. affinis* bat, but as SARS-CoV-2 spike can bind *R. affinis* ACE2 proteins comparably to RaTG13 spike binding (31), the divergence of the RaTG13 RBM is unlikely to be the result of optimization for *R. affinis* ACE2 sequences. The divergent RaTG13 RBM could reflect adaptation to an unsampled ACE2 or even optimization to allow generalist binding to multiple ACE2 sequences.

Alternatively, some of the RaTG13 specific RBD variation may reflect selective pressures unrelated to receptor binding, including escape from adaptive immunity (32). Interestingly, in the native bat host context, immune escape may not be a significant driver of spike evolution; for instance, BANAL coronavirus spikes are remarkably similar to the original SARS-CoV-2 spike despite much greater evolutionary distance from BANAL coronaviruses to SARS-CoV-2 than from SARS-CoV-2 to Omicron. Thus, if immune evasion played a role in RaTG13’s accumulation of spike variants, this could indicate important differences along the RaTG13 lineage, such as a different host or tissue tropism that led to greater pressure to escape adaptive immunity. The host, receptor environment, and evolutionary history of the lineage that contributed to the RaTG13 RBM are unknown and thus we can only speculate on the selective forces responsible for it. Isolation of more coronaviruses related to RaTG13 could greatly increase our understanding of this lineage.

## Materials and Methods

### Spike plasmid construction

We generated an initial set of spike-containing plasmids. Synthetic gene fragments (gblocks; IDT) were designed to encode the RBD residues 331-531 from RaTG13, with either arginine or threonine at residue 403 (SMG371, 372). Each fragment contained flanking sequences homologous to the regions adjacent to the RBD within the yeast surface display plasmid pETcon_SARS-CoV-2_RBD (Addgene 166782, deposited by Jesse Bloom) to enable Gibson assembly. This plasmid carries the SARS-CoV-2 RBD preceded by a galactose-inducible promoter and a methionine start codon, and followed by a stop codon and terminator. All sequences were yeast codon optimized to facilitate expression in *Saccharomyces cerevisiae*.

Gene fragments were amplified using KAPA HiFI HotStart DNA polymerase (Roche) and primers SMG383 and 384. The pETcon plasmid was digested with NdeI and XhoI, which removed the original RBD and linearized the plasmid, followed by gel purification. Inserts were assembled into the vector by Gibson Assembly at a 3:1 molar ratio of insert to vector and transformed into DH5alpha chemically competent cells. Transformants were selected on LB+AMP and plasmid DNA from individual colonies was screened by Sanger sequencing.

### Mosaic oligo library design

We developed a custom and easily modifiable Python script to generate the minimal number of oligonucleotide sequences for mosaic libraries by leveraging IUPAC degenerate bases. In this experiment, we focused on the receptor-binding motif (RBM, residues 439-505) and systematically generated all possible combinations of amino acid differences between the aligned RBM regions of RaTG13 and SARS-CoV-2.

To use this script, the user provides the two amino acid sequences of the region of interest and the desired flanking nucleotide sequences for cloning. The script aligns the two sequences, identifies all variable sites, and determines if each amino acid pair can be encoded by a single IUPAC degenerate codon (enabling a single oligonucleotide to encode both possible amino acids at that position) or if the two amino acids options must be ‘hardcoded’ using separate codons for each amino acid. The minimal set of amino acid sequences encoding the full combinatorial set of hardcoded differences is translated into nucleotide sequences using a yeast-specific codon usage table and IUPAC codes for the degenerate nucleotides. The script outputs a CSV file listing the full nucleotide sequences with the user defined flanking sequences. In addition, the script prints a summary table of all variable positions and if they are encoded with a degenerate or hardcoded codon.

We applied this script to the RBMs of SARS-CoV-2 and RaTG13. We provided flanking regions with 15 bases of homology to the RaTG13 RBM such that the cloned oligos would be in frame. Out of 17 variant sites, 12 could be coded with degenerate codons, while 5 needed to be hardcoded, and thus 2^5 = 32 oligonucleotides were required. The resulting nucleotide sequences were ordered from IDT as an oPool.

### Plasmid pool cloning

Linearized forms for the pETcon plasmids containing the RaTG13 RBD with R403 and T403 were generated by inverse PCR using Herculase II Fusion DNA polymerase (Agilent catalog # 600679) and primers (SMG393 and 394) that flanked the oPool insertion site. We gel purified (Qiagen catalog # 28074) the resulting linearized vectors.

The oPool was PCR amplified using KAPA HiFi HotStart DNA polymerase (Roche catalog # 9420398001) and primers complementary to the flanking constant regions (SMG379 and 380). The reverse primer contained 5 randomized nucleotides at 4-fold synonymous positions in flanking codons to introduce sequence barcodes. The amplified pool was gel purified. The amplified oPool was assembled into the vector by Gibson Assembly at a 5:1 molar ratio of insert to vector. Reactions were incubated at 50 C for 1 hour. We used the Clean and Concentrate kit (Zymo catalog # d4013) to concentrate the assembled plasmids. Lucigen Duo electrocompetent cells (Lucigen catalog # 60502-1) were electroporated using a MicroPulser Electroporator (Biorad catalog # 1652100). After a 1 hour outgrowth at 37 degrees, transformants were selected on LB+AMP plates.

By dilution plating, we estimated approximately 1.5 million colonies for each backbone (R403 and T403), providing greater than fivefold coverage of the theoretical library. Colonies were harvested by scraping, pooled by backbone, and grown overnight in LB+AMP, and plasmid DNA was purified by Qiagen Maxi Prep (catalog # 12963).

### PacBio sequencing of combinatorial variant libraries

The mosaic libraries were sequenced using PacBio circular consensus sequencing (CCS) to identify the genotype of each plasmid in the mosaic library and its associated synonymous barcode. Plasmid DNA was digested with AgeI and SacI to generate ∼1.8 kb linear fragments spanning the RBD and barcode region. Following a 2 hour digestion at 37 C, DNA was purified using a QIAquick PCR purification kit and size-selected using a SageELF instrument. The resulting DNA was then subjected to CCS HiFi sequencing on a Sequel II instrument.

CCS reads were processed using custom Python scripts to reconstruct the genotype of each mosaic construct and associate it with its synonymous barcode, generating a lookup table used for downstream genotype analysis.

### Yeast transformation

AWY101 (Wentz and Shusta, 2007; received from John Misasi), a strain designed for yeast surface display, was cultured at 30 C in selective media containing 6.7 g/L yeast nitrogen base, 5.0 g/L casamino acids, 1.065 g/L MES, and 2% w/v carbon source. Dextrose was used for routine maintenance and galactose supplemented with 0.1% dextrose was used to induce RBD expression. The plasmid libraries were transformed into AWY101 using a standard large-scale high efficiency lithium acetate transformation protocol (Gietz and Schiestl, 2007) and plated onto 15 cm selection media plates. Transformation efficiency was estimated by plating a 1:100 dilution, yielding approximately 400,000 colonies for each plasmid backbone. Colonies were harvested by washing plates and pooling for subsequent induction and binding assays.

### Yeast surface display assay

Yeast surface display and ACE2 binding measurements were performed using a protocol adapted from Starr et al. (11). Yeast libraries were grown overnight in selection media. 2x10^7 cells were washed once with media lacking a carbon source, and resuspended in 1.5 mL of media containing galactose and supplemented with 0.1% dextrose to induce surface expression of the RBD. Induction was carried out overnight (16-18 hrs) at room temperature overnight with continuous rotation.

Following induction, 0.67 OD600 of cells were collected for each binding reaction, washed twice with ice-cold PBS-BSA (0.2 mg/mL) and split into two reactions, one of which was incubated overnight at room temperature with 100 nM biotinylated monomeric human ACE2 protein (ACROBiosystems AC2-H82E8), and the other of which was not treated with ACE2.

Cells were subsequently washed twice with ice-cold PBS-BSA and fluorescently labeled for 1 hour at 4 C with agitation using PE-conjugated streptavidin (1:200; Thermo Fisher, S8666) to detect bound ACE2 and FITC-conjugated chicken anti-Myc (1:200; ICL CMYC-45F) to detect RBD surface expression.

Cells were washed twice and resuspended in ice-cold PBS-BSA and sorted via BD FACS Aria III. Gates were set based on singly labeled controls (PE or FITC only) and control strains (wild-type SARS-CoV-2 RBD or wild-type RaTG13 RBD in pETcon). We sorted 2 million single Myc+ cells into four bins based on the PE signal.

After an overnight recovery, plasmid DNA was extracted via a modified QIAprep spin MiniPrep protocol. Briefly, cultures were pelleted, suspended in 250 μL QIAprep P1 buffer and 2 μL zymolase (2.5 units per μL; MP Biomedicals catalog # 08320921) and incubated at 37 C for 30 minutes. QIAprep P2 buffer (250 μL) was then added and the plasmid purification was completed according to the manufacturer’s protocol.

### Amplicon sequencing

To quantify the abundance of each mosaic in each fluorescence bin, the RBD region was amplified from the plasmid DNA and sequenced using Illumina short read sequencing. Libraries were prepared using a two-step PCR protocol. The first PCR amplified the RBD and synonymous barcode region using Kapa HiFi HotStart ReadyMix and primers SMG423-432. To increase sequence diversity, forward and reverse primers consisted of pools of five primers containing 0-4 randomized nucleotides between the Nextera transposase adapter sequence and the gene-specific priming sequence. PCR products were purified using a MinElute PCR purification kit (Qiagen) and quantified with a Qubit dsDNA High Sensitivity Assay (Thermo Fisher).

A second PCR was used to attach Illumina adapters and Nextera dual indices for multiplexing. Briefly, 10 ng of purified amplicon DNA was amplified using Nextera adapter index primers and Kapa HiFi HotStart ReadyMix. Libraries were size selected and purified on a Size Select II E-gel (Invitrogen), quantified by Qubit dsDNA High Sensitivity kit, normalized to 4 nM, pooled, and sequenced on an Illumina Nextseq 2000 instrument following the manufacturer’s library preparation protocol.

### Data analysis

Illumina paired-end reads were merged using PEAR (33). Custom Python scripts were used to extract the RBD sequence and associated synonymous barcode from each read. Genotype-barcode pairs were filtered based on presence in the PacBio lookup table. Genotypes were filtered for those that were present in at least 2 fluorescence bins and had at least 10 reads total.

Relative ACE2 binding scores were calculated using custom R scripts. For each genotype, read counts within each fluorescence bin were normalized by the total number of sequencing reads and the number of cells sorted into that bin. The normalized bin frequencies were used to calculate a weighted average binding score, with higher values corresponding to stronger ACE2 binding. To visualize the effect of each variable position, mosaics were grouped according to whether they carried the SARS-CoV-2 or RaTG13 amino acid at that position. The distributions of ACE2 binding were plotted as violin plots. Statistical significance was determined from the coefficient for each position in an additive linear model containing all 18 positions. P values for the 18 positions’ coefficients were adjusted for multiple comparisons using the Benjamini-Hochberg false discovery rate (FDR). The position effect was calculated as the difference in mean bin score between mosaics carrying the SARS-CoV-2 and RaTG13 residues at each position. Positive values indicate that the SARS-CoV-2 residue increased ACE2 binding on average and negative values indicate that the RaTG13 residue had higher binding.

To determine how much phenotypic variation could be accounted for by different sets of positions, we fit linear models predicting mean bin score from the parental amino acid identities at the specified positions. Each position was encoded to whether it contained the SARS-CoV-2 or RaTG13 residue, and models contained only single position terms without interactions. Model fit was assessed using variance explained (adjusted R^2). We compared models containing six (449, 486, 493, 498, 501, and 505), seven (+403), eight (+439), and all 18 variable residues.

To examine context dependence at positions 493 and 498, each focal position was analyzed in combination with each of the other 17 variable positions. For each pair of positions, mosaics were divided into the four possible combinations of the SARS-CoV-2 and RaTG13 amino acids. We compared the effect of exchanging the parental residue at position 493 or 498 between mosaics carrying either parental residue at the second position. We fit a linear model containing the main effects and pairwise interactions. P values were adjusted as above. Because mean bin score is a nonlinear measure of ACE2 binding, we focused on interactions in which an effect on binding was reversed by another mutation. Positions 484 and 494 were identified as sites whose effects were flipped by 493 and 498. To examine their combined effects, mosaics were partitioned into all eight combinations of parental amino acids at positions 484, 494 and 493 or 498.

To determine how genotype sampling affected estimates of single position effects, we randomly downsampled the observed mosaic library across a range of library sizes. For each library size, 1000 replicates of genotypes were sampled and the effects of all 18 positions were recalculated as above. For each replicate, we calculated the root mean square deviation (RMSD) between the position effect estimates obtained from the downsampled library and the estimates from the complete library. To assess recovery of the major effect positions (403, 439, 486, 493, 498, 501, 505), we determined whether the sign of each effect in the downsampled library matched that estimated from the complete library. The fraction of replicates recovering the correct direction was determined at each library size.

We separately downsampled the library to determine how library size affected detection of the context dependence at positions 493 and 498. We considered mosaics in two sequence contexts: those containing the RaTG13 residues T484 and R494, and those containing the SARS-CoV-2 residues E484 and S494. Within each of these contexts, mosaics could carry either the RaTG13 or SARS-CoV-2 residue at 493 or 498, resulting in four genotypes for each analysis. At each downsampled library size, 1000 random replicate subsets were generated. For each replicate, we fit a model containing terms for 493 or 498, the sequence context, and their interaction. We asked if the effect of 493 or 498 differed between the RaTG13-like and SARS-CoV-2-like contexts. The fraction of downsamplings in which the interaction was statistically significant (p-value < 0.05) was used as the frequency with which context dependence was detected.

## Acknowledgments

We thank Tyler Starr and Michael Desai for resources and helpful discussion. We thank Luke Reding and all members of the Sadhu lab for helpful discussion. Next-generation sequencing was performed with the Microarrays and Single-Cell Genomics Core of the National Human Genome Research Institute. This work utilized the computational resources provided by the NIH HPC Biowulf Cluster (http://hpc.nih.gov). This work was supported by the Intramural Research Program of the National Human Genome Research Institute, NIH (1ZIAHG200401). The contributions of NIH authors are considered Works of the US Government. The findings and conclusions presented in this article are those of the authors and do not necessarily reflect the views of the NIH or the US Department of Health and Human Services.

## Artificial Intelligence (AI) Disclosure

We used ChatGPT (GPT-4 Sol) for assistance in generating code for oligo design and data analysis.

## Author Contributions

Conceptualization: S.M.G., M.J.S.; Investigation: S.M.G.; Methodology: S.M.G., M.J.S.; Writing: S.M.G., M.J.S.

## References

1. M. Hoffmann, et al., SARS-CoV-2 Cell Entry Depends on ACE2 and TMPRSS2 and Is Blocked by a Clinically Proven Protease Inhibitor. Cell 181, 271–280.e8 (2020).

2. B. Korber, et al., Tracking Changes in SARS-CoV-2 Spike: Evidence that D614G Increases Infectivity of the COVID-19 Virus. Cell 182, 812–827.e19 (2020).

3. J. A. Plante, et al., Spike mutation D614G alters SARS-CoV-2 fitness. Nature 592, 116–121 (2021).

4. P. Mlcochova, et al., SARS-CoV-2 B.1.617.2 Delta variant replication and immune evasion. Nature 599, 114–119 (2021).

5. P. Zhou, et al., A pneumonia outbreak associated with a new coronavirus of probable bat origin. Nature 579, 270–273 (2020).

6. M. Letko, A. Marzi, V. Munster, Functional assessment of cell entry and receptor usage for SARS-CoV-2 and other lineage B betacoronaviruses. Nat Microbiol 5, 562–569 (2020).

7. X.-Y. Ge, et al., Coexistence of multiple coronaviruses in several bat colonies in an abandoned mineshaft. Virol. Sin. 31, 31–40 (2016).

8. D. Cantoni, et al., Pseudotyped Bat Coronavirus RaTG13 is efficiently neutralised by convalescent sera from SARS-CoV-2 infected patients. Commun Biol 5, 409 (2022).

9. A. G. Wrobel, et al., SARS-CoV-2 and bat RaTG13 spike glycoprotein structures inform on virus evolution and furin-cleavage effects. Nat Struct Mol Biol 27, 763–767 (2020).

10. K. Liu, et al., Binding and molecular basis of the bat coronavirus RaTG13 virus to ACE2 in humans and other species. Cell 184, 3438–3451.e10 (2021).

11. T. N. Starr, et al., Deep Mutational Scanning of SARS-CoV-2 Receptor Binding Domain Reveals Constraints on Folding and ACE2 Binding. Cell 182, 1295–1310.e20 (2020).

12. F. Zech, et al., Spike residue 403 affects binding of coronavirus spikes to human ACE2. Nat Commun 12, 6855 (2021).

13. C. Conceicao, et al., The SARS-CoV-2 Spike protein has a broad tropism for mammalian ACE2 proteins. PLoS Biol 18, e3001016 (2020).

14. J. Zahradník, et al., SARS-CoV-2 variant prediction and antiviral drug design are enabled by RBD in vitro evolution. Nat Microbiol 6, 1188–1198 (2021).

15. T. N. Starr, et al., Deep mutational scans for ACE2 binding, RBD expression, and antibody escape in the SARS-CoV-2 Omicron BA.1 and BA.2 receptor-binding domains. PLoS Pathog 18, e1010951 (2022).

16. A. Moulana, et al., Compensatory epistasis maintains ACE2 affinity in SARS-CoV-2 Omicron BA.1. Nat Commun 13, 7011 (2022).

17. A. Kuo, C. Zhong, W. S. Lane, R. Derynck, Transmembrane transforming growth factor-α tethers to the PDZ domain-containing, Golgi membrane-associated protein p59/GRASP55. EMBO J 19, 6427–6439 (2000).

18. W. Ren, et al., Difference in Receptor Usage between Severe Acute Respiratory Syndrome (SARS) Coronavirus and SARS-Like Coronavirus of Bat Origin. J Virol 82, 1899–1907 (2008).

19. C. Liu, et al., HKU25 clade MERS-related coronaviruses use ACE2 as a functional receptor. Nat Microbiol 10, 2860–2874 (2025).

20. F. J. Poelwijk, M. Socolich, R. Ranganathan, Learning the pattern of epistasis linking genotype and phenotype in a protein. Nat Commun 10, 4213 (2019).

21. A. M. Phillips, et al., Binding affinity landscapes constrain the evolution of broadly neutralizing anti-influenza antibodies. eLife 10, e71393 (2021).

22. J. L. Tenthorey, et al., Indels allow antiviral proteins to evolve functional novelty inaccessible by missense mutations. Cell Genomics 5, 100818 (2025).

23. J. Lan, et al., Structure of the SARS-CoV-2 spike receptor-binding domain bound to the ACE2 receptor. Nature 581, 215–220 (2020).

24. E. C. Thomson, et al., Circulating SARS-CoV-2 spike N439K variants maintain fitness while evading antibody-mediated immunity. Cell 184, 1171–1187.e20 (2021).

25. E. Laurini, D. Marson, S. Aulic, A. Fermeglia, S. Pricl, Molecular rationale for SARS-CoV-2 spike circulating mutations able to escape bamlanivimab and etesevimab monoclonal antibodies. Sci Rep 11, 20274 (2021).

26. H. Guo, et al., Evolutionary Arms Race between Virus and Host Drives Genetic Diversity in Bat Severe Acute Respiratory Syndrome-Related Coronavirus Spike Genes. J Virol 94, e00902–20 (2020).

27. T. N. Starr, et al., ACE2 binding is an ancestral and evolvable trait of sarbecoviruses. Nature 603, 913–918 (2022).

28. M. F. Boni, et al., Evolutionary origins of the SARS-CoV-2 sarbecovirus lineage responsible for the COVID-19 pandemic. Nat Microbiol 5, 1408–1417 (2020).

29. H. Wang, L. Pipes, R. Nielsen, Synonymous mutations and the molecular evolution of SARS-CoV-2 origins. Virus Evol 7, veaa098 (2021).

30. P. Li, et al., Effect of polymorphism in Rhinolophus affinis ACE2 on entry of SARS-CoV-2 related bat coronaviruses. PLoS Pathog 19, e1011116 (2023).

31. S. Temmam, et al., Bat coronaviruses related to SARS-CoV-2 and infectious for human cells. Nature 604, 330–336 (2022).

32. F. P. Lyngse, et al., Household transmission of the SARS-CoV-2 Omicron variant in Denmark. Nat Commun 13, 5573 (2022).

33. J. Zhang, K. Kobert, T. Flouri, A. Stamatakis, PEAR: a fast and accurate Illumina Paired-End reAd mergeR. Bioinformatics 30, 614–620 (2014).

