## Supplementary Information for "Combinatorial mosaics dissect the residue level basis of ACE2 binding divergence between SARS-CoV-2 and RaTG13"

### Figures

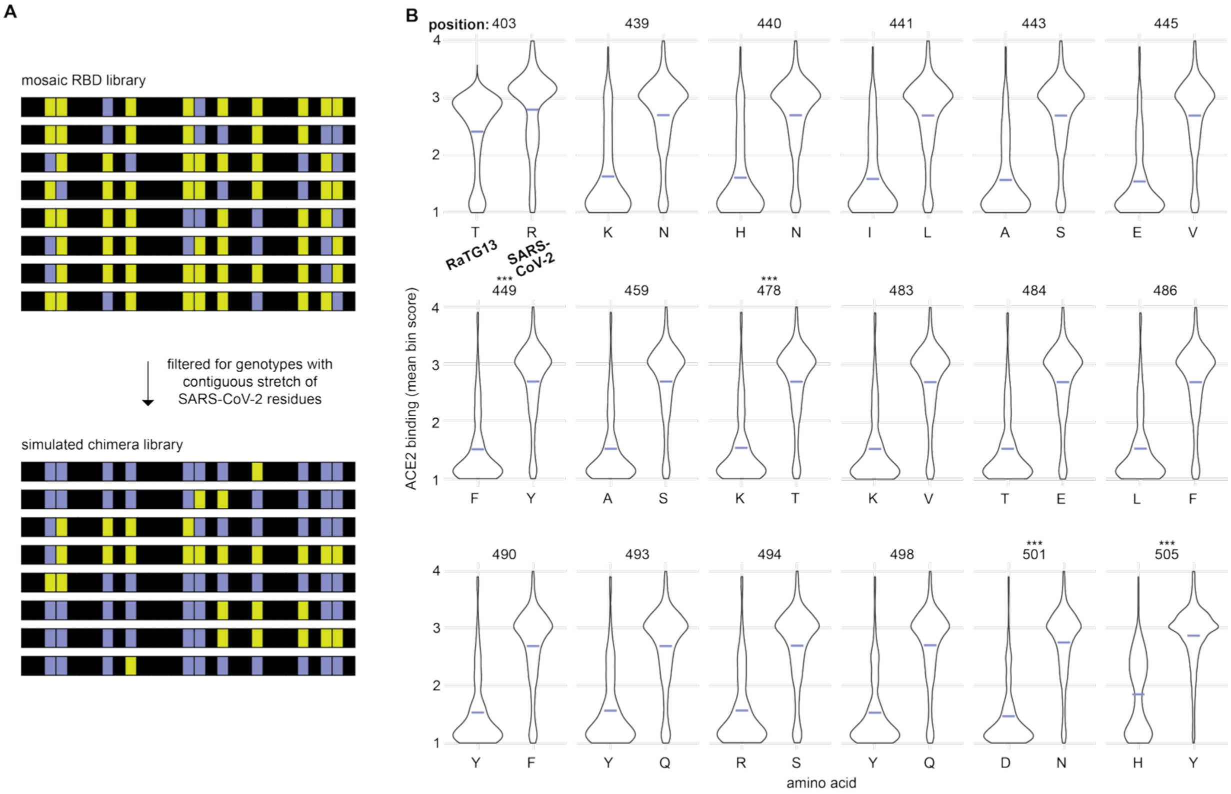


**Fig. S1. Simulated chimera analysis masks individual effects.** Genotypes from the mosaic library were restricted to those containing contiguous SARS-CoV-2 sequence to approximate traditional chimeras and assess how linkage affects apparent residue-level effects. **A.** Schematic of the simulated chimera analysis. Genotypes from the full mosaic library were restricted to those corresponding to chimeras, defined as containing a contiguous stretch of SARS-CoV-2 residues within an RaTG13 sequence. **B.** For each variable position, genotypes were separated according to whether they carried the RaTG13 or SARS-CoV-2 amino acid at that position. Violin plots show the distribution of mean bin scores. Purple dashes indicate the mean. Statistical significance was assessed with a linear model of ACE2-binding phenotypes fitted against the 18 variant sites in the simulated chimeras, and P values were adjusted for multiple comparisons using the Benjamini-Hochberg false discovery rate method (FDR < 0.05; adjusted p < 0.001 ***).

**Tables**

**Table S1. Strain Table**

| | |
|---|---|
| MSY84 | AWY101 AGA1::GAL1-AGA1::URA3 PDI1::GAPDH-PDI1::LEU2 ura3-52 trp1 leu2Δ1 his3Δ200 pep4::HIS3 prb1Δ1.6R can1 GAL |
| MSY334 | AWY101 pETCON - WT RaTG13 RBD |
| MSY335 | AWY101 pETCON - T403R RaTG13 RBD |
| MSY286 | AWY101 AWY101 RaTG13/SARS-CoV-2 RBD mosaic pool in WT RaTG13 (T403) |
| MSY287 | AWY101 RaTG13/SARS-CoV-2 RBD mosaic pool in MUT RaTG13 (R403) |

**Table S2. Oligonucleotide Table**

| | | |
|---|---|---|
| SMG371 | GAGGGTCGGCTAGCCATATGAACATAACTAA<br>CTTATGTCCATTTGGGGAAGTGTTCAACGCTA<br>CGACATTTGCCAGCGTGTATGCATGGAACAG<br>AAAGAGAATTTCAAATTGCGTCGCTGACTATT<br>CAGTCTTGTATAACAGCACGTCCTTTTCAACC<br>TTCAAGTGTTACGGTGTCAGTCCAACTAAATT<br>AAATGATCTATGTTTCACTAACGTATATGCAG<br>ACAGCTTTGTTATTACCGGAGACGAGGTGAG<br>GCAGATCGCGCCCGGTCAGACTGGAAAGATT<br>GCGGATTACAACTACAAGCTACCAGACGACT<br>TCACTGGATGTGTAATCGCCTGGAACTCCAA<br>GCATATCGACGCAAAAGAGGGCGGGAACTTT<br>AATTACTTATACCGTCTATTCAGAAAAGCAAA<br>CTTAAAACCTTTTGAGAGGGACATTTCTACAG<br>AAATATATCAAGCCGGGTCAAAACCGTGCAA<br>CGGACAAACCGGGCTTAATTGCTATTATCCA<br>CTGTATAGGTATGGCTTTTATCCTACTGATGG<br>CGTAGGGCACCAGCCCTACAGGGTTGTTGTT<br>TTATCCTTCGAGCTTTTGAATGCCCCGGCGA<br>CCGTGTGCGGGCCCAAGAAGTCTACTCTCGA<br>GGGGGGCGGTTCCGA | Yeast codon optimized wild-type RaTG13 RBD (amino acids 331-551), ordered as a gene block from IDT |
| SMG372 | GAGGGTCGGCTAGCCATATGAACATAACTAA<br>CTTATGTCCATTTGGGGAAGTGTTCAACGCTA<br>CGACATTTGCCAGCGTGTATGCATGGAACAG<br>AAAGAGAATTTCAAATTGCGTCGCTGACTATT<br>CAGTCTTGTATAACAGCACGTCCTTTTCAACC<br>TTCAAGTGTTACGGTGTCAGTCCAACTAAATT<br>AAATGATCTATGTTTCACTAACGTATATGCAG<br>ACAGCTTTGTTATTAGAGGAGACGAGGTGAG<br>GCAGATCGCGCCCGGTCAGACTGGAAAGATT<br>GCGGATTACAACTACAAGCTACCAGACGACT<br>TCACTGGATGTGTAATCGCCTGGAACTCCAA<br>GCATATCGACGCAAAAGAGGGCGGGAACTTT<br>AATTACTTATACCGTCTATTCAGAAAAGCAAA<br>CTTAAAACCTTTTGAGAGGGACATTTCTACAG<br>AAATATATCAAGCCGGGTCAAAACCGTGCAA<br>CGGACAAACCGGGCTTAATTGCTATTATCCA<br>CTGTATAGGTATGGCTTTTATCCTACTGATGG<br>CGTAGGGCACCAGCCCTACAGGGTTGTTGTT<br>TTATCCTTCGAGCTTTTGAATGCCCCGGCGA<br>CCGTGTGCGGGCCCAAGAAGTCTACTCTCGA<br>GGGGGGCGGTTCCGA | Yeast codon optimized T403R RaTG13 RBD (amino acids 331-551), ordered as a gene block from IDT |
| SMG375 | CATATGGCTAGCCGACCCTC | Amplifies pETCON plasmid for insertion of RBD gene blocks |
| SMG376 | CTCGAGGGGGGCGGTTCCGA | Amplifies pETCON plasmid for insertion of RBD gene blocks |

| | | |
|---|---|---|
| SMG383 | GAGGGTCGGCTAGCCATATG | Amplifies RBD gene blocks for insertion into linearized pETCON |
| SMG384 | TCGGAACCGCCCCCCTCGAG | Amplifies RBD gene blocks for insertion into linearized pETCON |
| SMG379 | AAGCTACCAGACGACTTCACTGGATGTGTAATCGCCTGGAACTCC | Amplifies IDT oligo pool for insertion into RaTG13 RBD in pETCON |
| SMG380 | GCACACGGTCGCCGGGGCATTCAAAAGCTCGAANGANAGNACNACNACCCTGTAGGGCTG | Amplifies IDT oligo pool for insertion in RaTG13 RBD in pETCON |
| SMG393 | CAGTGAAGTCGTCTGGTAGCTTG | Amplifies pETCON - T403 and R403 RaTG13 RBD for insertion of oligo pool |
| SMG394 | GCCCCGGCGACCGTGTGCGGGCCCAAGAAGTCTACTC | Amplifies pETCON - T403 and R403 RaTG13 RBD for insertion of oligo pool |
| SMG423 | TCGTCGGCAGCGTCAGATGTGTATAAGAGACAGCAAGCTACCAGACGACTTCACTG | Amplifies RBD mosaic region for NGS |
| SMG424 | TCGTCGGCAGCGTCAGATGTGTATAAGAGACAGNCAAGCTACCAGACGACTTCACTG | Amplifies RBD mosaic region for NGS |
| SMG425 | TCGTCGGCAGCGTCAGATGTGTATAAGAGACAGNNCAAGCTACCAGACGACTTCACTG | Amplifies RBD mosaic region for NGS |
| SMG426 | TCGTCGGCAGCGTCAGATGTGTATAAGAGACAGNNNCAAGCTACCAGACGACTTCACTG | Amplifies RBD mosaic region for NGS |
| SMG427 | TCGTCGGCAGCGTCAGATGTGTATAAGAGACAGNNNNCAAGCTACCAGACGACTTCACTG | Amplifies RBD mosaic region for NGS |
| SMG428 | GTCTCGTGGGCTCGGAGATGTGTATAAGAGACAGGATCTCTATTACAAGTCCTCTTCAG | Amplifies RBD mosaic region for NGS |
| SMG429 | GTCTCGTGGGCTCGGAGATGTGTATAAGAGACAGNGATCTCTATTACAAGTCCTCTTCAG | Amplifies RBD mosaic region for NGS |
| SMG430 | GTCTCGTGGGCTCGGAGATGTGTATAAGAGACAGNNGATCTCTATTACAAGTCCTCTTCAG | Amplifies RBD mosaic region for NGS |
| SMG431 | GTCTCGTGGGCTCGGAGATGTGTATAAGAGACAGNNNGATCTCTATTACAAGTCCTCTTCAG | Amplifies RBD mosaic region for NGS |
| SMG432 | GTCTCGTGGGCTCGGAGATGTGTATAAGAGACAGNNNNGATCTCTATTACAAGTCCTCTTCAG | Amplifies RBD mosaic region for NGS |
